# Interferon lambda drives immunological maturation in the infant lung and protects against lethal *Bordetella pertussis* infection

**DOI:** 10.64898/2026.09.22.753444

**Authors:** Da’Kuawn Johnson, Nicholas H. Carbonetti

## Abstract

Serious pertussis infections disproportionately affect infants but the biological basis for this age-dependent susceptibility remains unclear. Infant mouse models recapitulate features of severe human infant pertussis. We investigated the role of interferon lambda (IFN-λ), a key regulator of mucosal immunity, in *Bordetella pertussis* infection of infant mice. While infected adult mice upregulate lung IFN-λ, infant mice inoculated at P7 fail to upregulate IFN-λ and succumb to infection. We hypothesized that failure to produce IFN-λ during infection represents a critical immunological deficit in infant mice, and that restoring IFN-λ signaling would improve survival outcomes. Whereas wild-type mice gained complete protection from lethal *B. pertussis* infection by P10, mice lacking the IFN-λ receptor component IFNLR1 did not achieve full protection until P21. Loss of IFNLR1 was associated with enhanced bacterial dissemination to systemic organs. Infant mice possessed a functional IFN-λ receptor in the lungs but failed to upregulate IFN-λ during infection, and supplementing IFN-λ exogenously extended survival. RNA sequencing of lung tissue from infected and uninfected wild-type and IFNLR1-deficient mice inoculated at different ages identified an immune transcriptional framework distinguishing susceptible from resistant animals at a systems level and revealed IFN-λ signaling as a critical driver of immunological maturation in the infant lung. Infected infant IFNLR1-deficient mice had dysregulated immune cell recruitment to the lungs, indicating a quantitatively expanded but qualitatively impaired response. These findings demonstrate that IFN-λ affects immune maturation accounting for a critical window of age-dependent resistance to lethal pertussis with novel therapeutic possibilities for human infants with this disease.

## INTRODUCTION

*Bordetella pertussis*, the causative agent of whooping cough, has re-emerged as a significant public health threat following a lull in cases during the COVID-19 pandemic. In 2024, the United States reported the highest number of pertussis cases since 2012, with case counts more than six-fold higher than in 2023, and surveillance data through 2025 indicate that case counts remain elevated above pre-pandemic levels^1^.

Pertussis disproportionately affects the youngest individuals. Infants under six months of age, who have not yet completed their primary vaccination series, account for the majority of pertussis-related hospitalizations and deaths^1,2^. Approximately one third of infected infants under six months required hospitalization in 2024, and 57% of all pertussis-related deaths reported to the CDC from 2013 to 2022 occurred among infants under two months of age^1^. By contrast, older children and adults typically experience milder, self-limiting disease. Despite renewed clinical urgency, no consistently effective therapies exist for individuals suffering from severe pertussis, and the biological basis for this striking age gradient in disease severity remains incompletely understood.

Infant mouse models recapitulate several key features of severe human infant pertussis and have been instrumental in dissecting the immunological determinants of age-dependent susceptibility^3,4^. Infant mice inoculated at early postnatal ages develop lethal infection with increased pulmonary bacterial burden, systemic dissemination of bacteria to other organs, leukocytosis, and pulmonary hypertension, similar to the severe disease observed in hospitalized human infants^3,4^. Adult mice, by contrast, restrict infection to the respiratory tract, mount effective inflammatory responses, and recover^3–5^. Age at the time of inoculation is a critical determinant of outcome in this model, with a transition from susceptibility to resistance to lethal infection occurring across the first 2 postnatal weeks^4^.

The heightened susceptibility of neonates and young infants to certain respiratory pathogens reflects fundamental differences in immune ontogeny relative to older individuals^5–7^. Neonatal innate immune responses are broadly characterized by impaired pattern recognition receptor signaling, reduced cytokine production, and deficient effector cell function^5,6,8^. These differences are not merely quantitative — neonatal immune cells exhibit qualitatively distinct epigenetic and chromatin accessibility landscapes that contribute to altered activation programs^9,10^, potentially biasing responses toward tolerance rather than effective defense^6,8^. Despite these well-characterized general features of neonatal immunity, the specific immunological mechanisms that render young infants uniquely vulnerable to severe pertussis remain poorly defined.

We have studied the role of interferons (IFNs) in outcomes of *B. pertussis* infection in mouse models. IFNs are a family of cytokines originally defined by their ability to interfere with viral replication, and their antiviral functions have been extensively characterized across multiple tissues and infection models^11,12^. Beyond antiviral immunity, IFNs also exert context-dependent effects during bacterial infections^13–15^, where they can either enhance host defense or contribute to immune-mediated pathology. The IFN signaling network is composed of three major classes — type I (IFN-α/β), type II (IFN-γ), and type III (IFN-λ) IFNs — each defined by distinct cytokines, receptor complexes, and cellular targets. IFN-λ signals through a heterodimeric receptor composed of IFN lambda receptor 1 (IFNLR1) and interleukin-10 receptor beta (IL10RB)^16,17^. Unlike type I IFN receptors, which are broadly expressed^18^, IFNLR1 expression is largely restricted to epithelial cells at barrier surfaces^19–21^, including the respiratory tract^22–24^, and subsets of immune cells^25^. This restricted receptor distribution positions IFN-λ as a key regulator of mucosal immunity and epithelial homeostasis^26,27^. Notably, IL-22, another cytokine that shares the IL-10RB receptor subunit with IFN-λ, promotes epithelial barrier integrity and antimicrobial peptide production at mucosal surfaces^28^. IFN-λ has diverse effects in experimental bacterial infection models, being protective in some contexts and deleterious in others^13,15^.

Prior work from our laboratory demonstrated age-dependent roles for both type I and type III IFNs in the pathogenesis of *B. pertussis* infection in mouse models^5^. Infected adult mice upregulate expression of type I, type II, and type III IFNs in the lung, whereas infected infant mice inoculated at postnatal day 7 (P7) fail to upregulate these cytokines^4,5^. IFN-γ and natural killer (NK) cells, major early producers of IFN-γ, play protective roles against *B. pertussis* infection in both adult and infant mice^4,29,30^, but the roles and effects of type I and type III IFNs are age-dependent^5^. Since infected adult mice robustly upregulate IFN-λ in the lungs, while infected P7 mice fail to upregulate IFN-λ and succumb to infection^5^, we hypothesized that failure to produce sufficient IFN-λ during infection represents a critical immunological deficit in infant mice, and that restoring IFN-λ signaling would improve survival outcomes. To test this, we investigated the role of IFN-λ signaling across postnatal ages using an IFNLR1-deficient mouse model, characterized the IFN-λ production defect in infant mice, and performed bulk RNA sequencing from lung tissue to define the transcriptional programs regulated by IFN-λ signaling during infection. By comparing wild-type (WT) and IFNLR1-deficient mice under infected and uninfected conditions across postnatal ages, we identify an immune transcriptional framework that distinguishes susceptible from resistant animals at a systems level and reveals IFN-λ signaling as a critical driver of immunological maturation in the neonatal lung. Understanding these mechanisms has implications not only for pertussis — where improved neonatal vaccination strategies and adjunctive therapies are urgently needed — but also for the broader question of how IFN biology shapes respiratory immunity across postnatal development, with potential relevance to other respiratory pathogens that disproportionately affect young infants^31–33^.

## MATERIALS AND METHODS

### Mouse Strains

C57BL/6 mice were purchased from Charles River Laboratories or the Jackson Laboratory for in-house breeding. All mouse strains were on the C57BL/6 genetic background. IFNLR1^fl/fl^ and IFNLR1^−/−^ (hereafter referred to as IFNLR1 KO) were provided by S. Kotenko (Rutgers New Jersey Medical School)^24^. C57BL/6 mice were used as age-matched controls unless otherwise indicated. All studies were performed on pre-weaning animals aged postnatal day 7 (P7) to P21 or adult mice 6 to 8 weeks old. Experimental pups were obtained by timed mating in-house. All animal procedures were performed under protocols approved by the Institutional Animal Care and Use Committee at the University of Maryland School of Medicine. All mouse strains were housed, bred, and maintained at the animal facilities at the University of Maryland School of Medicine in accordance with institutional guidelines.

### Bacterial Infections

Bacteria were suspended in phosphate-buffered saline (PBS) and administered via aerosol by nebulizer (Pari Vios) for 20 min as previously described^3^. The resulting dose in a mouse 24 h post-inoculation is between 5 × 10^5^ colony-forming units (CFU)/lung and 2 × 10^6^ CFU/lung depending on the age of the mouse at the time of inoculation. Control mice were sham inoculated by aerosolized PBS. On the indicated days post-inoculation (dpi), lungs and other organs were removed to analyze bacterial burden, transcript levels, or protein levels. For survival studies, mice were observed over 21 dpi when the experiment was terminated, as previously described^4^. Generally, groups of adult mice consisted of 4 to 6 animals of both sexes and groups of infant mice were litters of 5 to 9 animals of both sexes.

### RNA Isolation and Library Preparation

Whole lungs were harvested at the indicated time points, homogenized, and total RNA was isolated using standard trizol-chloroform–based extraction. RNA-seq library preparation and sequencing were performed separately for the adult and infant mouse cohorts used in this study, corresponding to two batches collected in different years. Adult mouse samples were collected and sequenced by Maryland Genomics, Institute for Genome Sciences, University of Maryland School of Medicine, generating paired-end Illumina reads and targeting 50 million reads per sample. For infant mouse samples (P7 and P10), collected, mRNA was enriched using the NEBNext Poly(A) mRNA Magnetic Isolation Module (New England Biolabs, Ipswich, MA), and strand-specific, dual uniquely indexed libraries were generated using the NEBNext Ultra II Directional RNA Library Prep Kit for Illumina (New England Biolabs, Ipswich, MA). Infant libraries were sequenced on an Illumina NovaSeq 6000 using 100 bp paired-end reads, targeting 30 million read pairs per sample.

### RNAseq analyses and pipelines

RNA sequencing analysis was carried out by Maryland Genomics, Institute for Genome Sciences, UMSOM. Paired-end Illumina reads were mapped to the mouse reference genome (Ensembl release GRCm39.110) using HISAT2 (v2.1.0) with default mismatch parameters, and alignment statistics were generated using SAMtools (v1.17). Alignment metrics were comparable across experimental groups, indicating uniform sequencing depth and alignment performance.

Gene-level read counts for each annotated gene were calculated using HTSeq^34^. Raw counts were normalized using DESeq2 (v1.5.24)^35^-based size factor estimation to account for differences in library depth. Exploratory analyses, including boxplots of normalized counts, density plots, and principal component analysis (PCA), were used to assess sample-to-sample variability and overall clustering behavior. Samples clustered primarily by experimental condition, with no evidence of extreme outliers or batch effects.

Differential gene expression analyses were performed using DESeq2^35^. Genes were considered significantly differentially expressed using a false discovery rate (FDR) cutoff of ≤ 0.05 and an absolute log2 fold-change threshold of ≥ 1. Pairwise comparisons were performed between infected and control groups within genotype, as well as between genotypes under matched conditions. Summary statistics, MA plots, and heatmaps were used to visualize differential expression patterns. Complete DEG lists for all pairwise comparisons, including gene symbol, log2 fold change, p-value, and adjusted p-value, are provided in a supplemental file (Table S1).

For downstream analyses, normalized expression matrices and differential expression outputs were further processed using open-source R-based pipelines developed in-house. All custom analysis code will be deposited in a public repository (GitHub/Zenodo — accession pending). Gene-set enrichment analyses were performed using curated databases including Gene Ontology (GO)^36,37]^, Kyoto Encyclopedia of Genes and Genomes (KEGG)^38^, and Reactome^39^ using the clusterProfiler package^40^ with Benjamini-Hochberg correction. IPA upstream regulator analysis was performed using Ingenuity Pathway Analysis software (Qiagen)^41^.

Immune cell deconvolution was performed on normalized bulk RNA-seq expression matrices using two independent computational methods. mMCP-counter (murine Microenvironment Cell Population counter) estimates the abundance of 16 immune and stromal cell populations using highly specific transcriptomic marker gene sets derived from the ImmGen database and validated against flow cytometry data from multiple murine tissues^42^. seqImmuCC estimates immune cell composition using a cell-centric gene signature approach trained on murine RNA-seq data and was used as an independent corroborating method^43^. Deconvolution scores were compared across age and genotype groups and are reported as exploratory findings given the sample size of n=3 biological replicates per group.

### Flow Cytometry

Lungs were perfused by injection through the right ventricle with 10 mL PBS for mice aged ≤P21 or 20 mL PBS for adult mice to remove circulating leukocytes. Single-cell suspensions were generated by incubation of minced lung tissue in digestion buffer containing 2 mg/mL Collagenase Type XI (Sigma-Aldrich), 20 mg/mL DNase I (Roche), 5 mg/mL Liberase TM (Sigma-Aldrich), and 100 mg/mL Hyaluronidase type IS (Sigma-Aldrich) for 45 min at 37°C. Tissue was ground through a 100-μm cell strainer, leukocytes were separated on a 37% Percoll density gradient (Sigma-Aldrich), and red blood cells were removed by ACK lysis. Single-cell suspensions were counted by trypan blue staining.

A total of 2 × 10⁶ cells were stained with Zombie Aqua fixable viability dye (BioLegend) in PBS for 15 min on ice. Cells were washed twice in FACS buffer (PBS, 0.5% BSA, 2 mM EDTA) and incubated with Fc Block (BioLegend) for 15 min on ice. Cells were stained with the following fluorochrome-conjugated antibodies at 4°C for 15 min: anti-CD45 (clone 30-F11, BUV395, eBioscience/Invitrogen), anti-CD11b (clone M1/70, BV711, eBioscience/Invitrogen), anti-Ly6G (clone 1A8, BV421, eBioscience/Invitrogen), anti-CD3ε (clone 17A2, Alexa Fluor 700, BioLegend), and anti-NK1.1 (clone PK136, PE/Cyanine5, BioLegend). After staining, cells were fixed in 2% paraformaldehyde for 10 min at room temperature, washed, and resuspended in FACS buffer.

Cells were acquired on the Cytek Aurora spectral flow cytometer and analyzed using FlowJo v10 (BD). Absolute cell counts were determined using CountBright counting beads (Thermo Fisher). Fluorescence-minus-one controls were used to set gates for each antibody. Neutrophils were defined as CD45+CD11b+Ly6G+. NK cells were defined as CD45+CD3ε−NK1.1+. T cells were defined as CD45+CD3ε+.

### Statistical Analysis

Data analysis and graph generation were performed using GraphPad Prism version 11.0 (GraphPad Software) and R (version 4.3.1). Results are presented as mean ± standard deviation (SD). Statistical significance between two groups was assessed using an unpaired Student’s t-test. For comparisons among multiple groups, one-way analysis of variance followed by Tukey’s multiple comparisons test was used. Differential gene expression analysis was performed using the limma-voom framework^44^ as described above. A P value of < 0.05 was considered statistically significant.

## RESULTS

### IFN-λ signaling confers age-dependent protection against B. pertussis infection

We previously demonstrated that wild-type (WT) mice acquire age-dependent resistance to lethal *B. pertussis* infection, with mice inoculated at postnatal day 7 (P7) succumbing to infection, while mice inoculated at postnatal day 10 (P10) and older survive^4^. The transition between P7 and P10 therefore represents a critical window in which protective immunity becomes established (Fig 1A). To determine whether IFN-λ signaling contributes to this protection, we inoculated IFNLR1 KO mice at multiple ages and monitored survival. Unlike WT mice, which were fully protected by P10, the majority (80%) of IFNLR1 KO mice inoculated at P10 did not survive infection, and complete survival was not observed in IFNLR1 KO animals until P21 (Fig. 1B, p<0.0001). These data indicate that IFN-λ signaling is a major contributor to age-dependent protection between P10 and P21.

**Figure 1.**
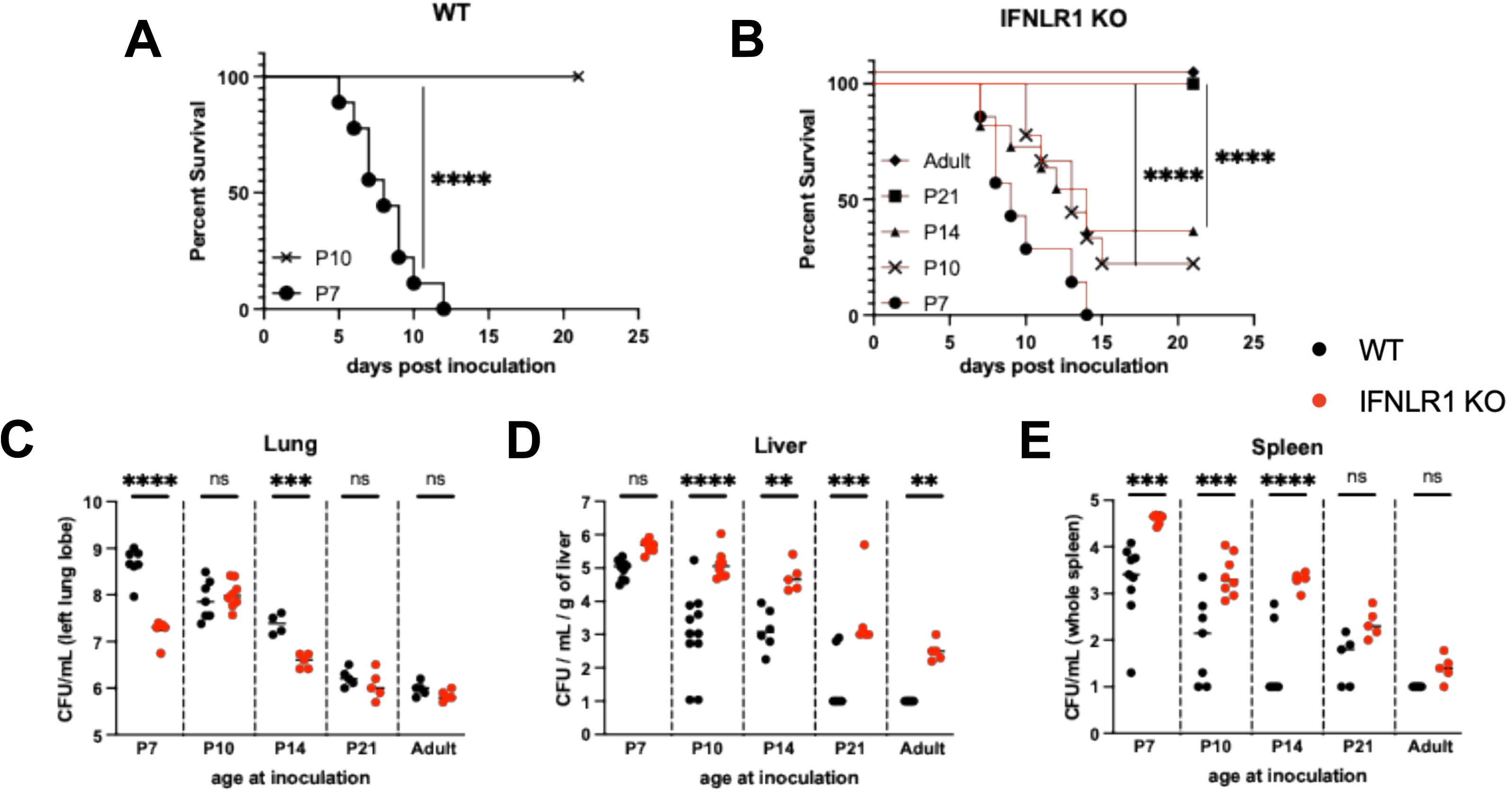
Young IFNLR1 KO mice exhibit enhanced susceptibility to *B. pertussis* infection. (A) Survival of wild-type (WT) mice inoculated with *B. pertussis* at P7 and P10 (n≥8 per group). (B) Survival of IFNLR1 KO mice inoculated at P7, P10, P14, P21, or adult age (6 wks). Statistical significance for survival experiments was determined by log-rank test (****p<0.0001; n≥8 per group). (C–E) Bacterial burden in lung (C), liver (D), and spleen (E) at day 7 post-inoculation in WT (black circles) and IFNLR1 KO (red circles) mice inoculated at the indicated ages. Each dot represents an individual animal. Statistical comparisons between WT and IFNLR1 KO within each age group were performed by lognormal one-way ANOVA with Holm-Šídák multiple comparisons test (ns, not significant; **p<0.01; ***p<0.001; ****p<0.0001; n≥5 per group).

Given this observed survival defect, we asked whether IFNLR1 KO mice also exhibited differences in bacterial burden and dissemination. We harvested lungs, liver and spleen at 7 days post-inoculation (dpi) from mice inoculated at the various ages and quantified bacterial burdens. Lung bacterial burdens were significantly higher in WT mice compared to IFNLR1 KO mice at P7 and P14, though differences were not significant at P10, P21, or in adults (Fig. 1C). However, IFNLR1 KO mice exhibited enhanced systemic dissemination - liver and spleen bacterial burdens were greater in IFNLR1 KO mice compared to WT controls across all ages, though differences did not reach statistical significance in all comparisons (Fig. 1D-E). Taken together, these data demonstrate that IFN-λ signaling limits systemic dissemination of *B. pertussis*, particularly in young mice.

Having established that IFN-λ signaling is required for age-dependent protection, we next asked whether a defect in the IFN-λ signaling axis may contribute to the susceptibility of P7 WT mice to lethal infection. To determine whether IFNLR1 is expressed and functional in infant lungs, we administered IFN-λ or IFN-β (as a positive control) intradermally to uninfected P7 and P10 WT mice and then probed lung homogenates for phosphorylated STAT1, since type I and III IFNs both signal through JAK-mediated STAT1 phosphorylation^16,17,45^. Both ages demonstrated robust STAT1 phosphorylation in response to both cytokines (Fig. 2A), confirming that IFNLR1 (as well as the type I IFN receptor) is expressed and functionally coupled to downstream signaling in the lungs of P7 mice, and that lack of a functional IFN-λ receptor does not account for the susceptibility of P7 mice. The comparatively stronger STAT1 phosphorylation observed with IFN-β (Fig. 2A) is consistent with the broader cellular distribution of the type I IFN receptor relative to the more restricted IFN-λ receptor .

**Figure 2.**
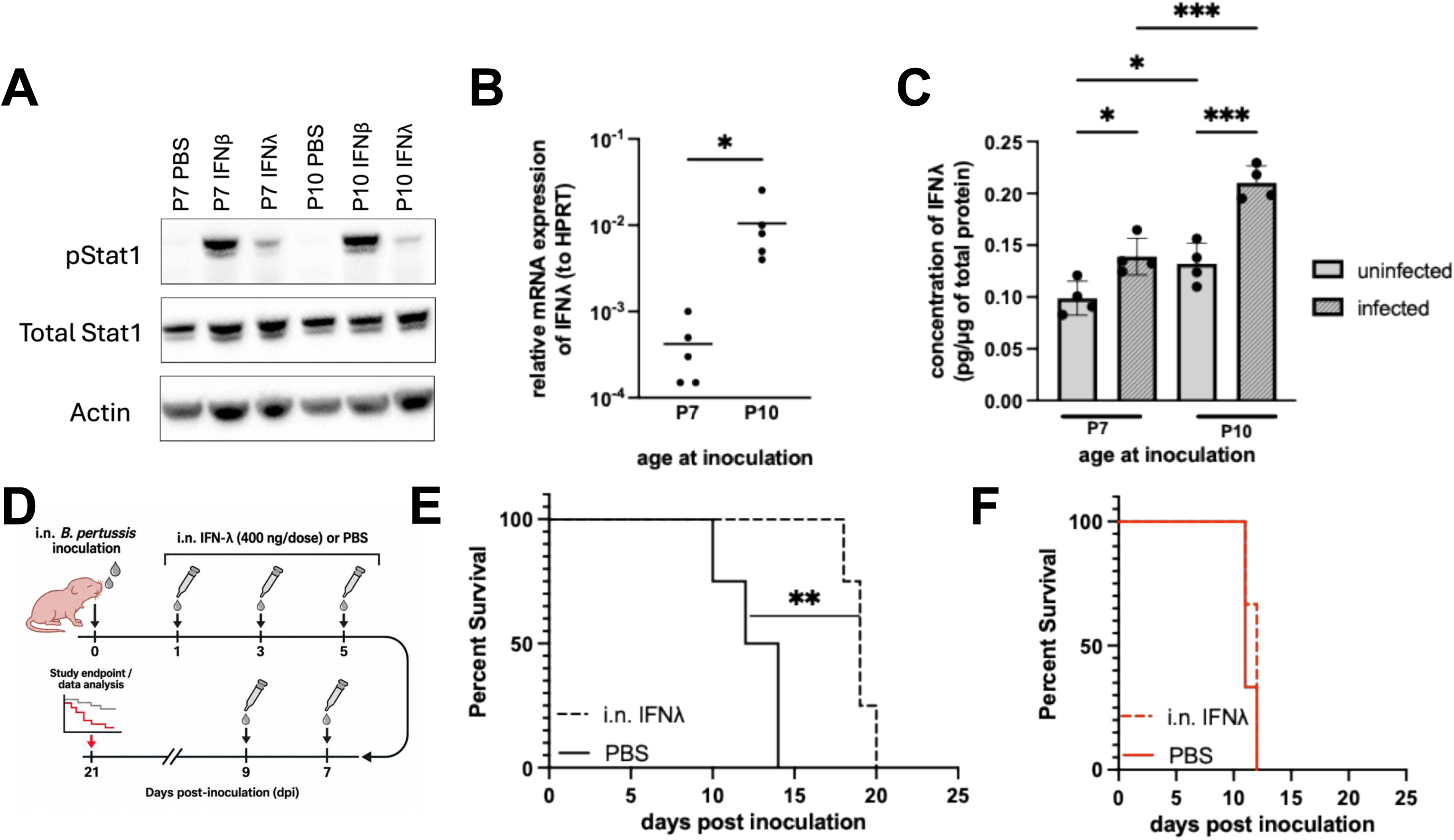
P7 WT mice express functional IFN-λ receptor but produce insufficient IFN-λ during *B. pertussis* infection. (A) Western blot analysis of phosphorylated STAT1 (pSTAT1), total STAT1, and β-actin in lung homogenates from P7 and P10 WT mice treated intradermally with PBS vehicle, 1 μg IFN-β, or 1 μg IFN-λ for 30 min. (B) Relative IFN-λ2/3 mRNA expression normalized to HPRT in lung homogenates at 7 dpi from WT mice inoculated at P7 or P10. Each dot represents an individual animal. Statistical comparison by unpaired two-tailed t-test (p=0.0326; n=5 per group). (C) IFN-λ2/3 protein concentration (pg/μg total protein) at 10 dpi in total lung homogenates from uninfected (light gray) and infected (dark gray with stripes) WT mice inoculated at P7 or P10. Statistical comparisons by ordinary one-way ANOVA with Holm-Šídák multiple comparisons test (*p<0.05; ***p<0.001; n=4 per group). (D) Schematic of the intranasal IFN-λ treatment regimen (i.n. IFN-λ; 400 ng/dose at 1, 3, 5, 7, and 9 dpi) or PBS control administered to infected P7 mice. (E) Survival of infected WT mice inoculated at P7 and treated intranasally with recombinant IFN-λ or PBS control (as in D). Statistical comparison by log-rank test (p=0.0091; n≥8 per group). (F) Survival of IFNLR1 KO mice inoculated at P7 and treated intranasally with recombinant IFN-λ or PBS control (as in D). n≥6 per group.

With receptor functionality established, we next examined whether infant mice express IFN-λ during infection. IFN-λ transcript levels and protein concentrations in lung homogenates from infected WT mice revealed a striking age-dependent difference. Infected P10 WT mice produced significantly higher levels of IFN-λ mRNA in the lungs compared to infected P7 WT mice (Fig. 2B, p=0.0326). IFN-λ protein concentrations were significantly elevated in infected P10 mice relative to both uninfected P10 controls and infected P7 mice (Fig. 2C). Infected P7 mice failed to mount a comparable induction above baseline (Fig. 2C). These data indicate that P7 animals fail to produce sufficient IFN-λ cytokine during infection despite having a functional receptor, suggesting that the inability to induce an adequate IFN-λ response, rather than any defect in the receptor itself, underlies their susceptibility. To directly test this, we treated infected P7 WT mice with intranasal purified IFN-λ and monitored survival (Fig. 2D). IFN-λ treatment significantly prolonged survival compared to PBS-treated WT controls (Fig. 2E, p=0.0091), demonstrating that augmenting IFN-λ signaling is sufficient to delay disease progression in susceptible infants. This effect was IFNLR1-dependent, as IFN-λ treatment failed to prolong survival in IFNLR1 KO animals (Fig. 2F), ruling out off-target effects of the cytokine treatment.

### IFN-λ signaling contributes to maturation of the infant lung transcriptional program

To gain mechanistic insight into how IFN-λ signaling shapes pulmonary responses to *B. pertussis* infection across age, we performed bulk RNA sequencing on whole lung tissue from WT and IFNLR1 KO mice infected with *B. pertussis* or sham-inoculated with PBS. Mice were inoculated at P7, P10, or adult age (6-8 weeks) and lungs were harvested at 7 dpi, yielding a total of 12 experimental groups that varied across 3 biological dimensions: age at inoculation, infection status, and IFNLR1 genotype. This design enabled us to dissect transcriptional responses to infection attributable to age, IFN-λ signaling, and their interaction.

To assess the overall structure of transcriptional variation across the dataset, we performed principal component analysis (PCA) on normalized gene expression values (Fig. S1A). Rather than segregating cleanly along a single principal component in PC1 vs PC2 analysis, samples separated along two diagonal axes that jointly reflected age and infection status (Fig. 3A–B). Adult samples occupied the upper right of the PCA plot, while P7 and P10 samples occupied the lower left, indicating that age contributes to both PC1 and PC2 rather than a single axis. Along the orthogonal diagonal, PBS-inoculated samples clustered toward the upper left and *B. pertussis*-infected samples toward the lower right, indicating that infection status similarly contributes to both components. Inspection of the genes contributing most strongly to each principal component revealed enrichment for inflammatory genes in both PC1 and PC2 (Table S2 and S3), consistent with age and infection status each shaping overlapping, rather than independent, components of transcriptional variance in the lung. Analysis of PC3 in contrast revealed a distinct contribution of mouse genotype (WT vs IFNLR1 KO) (Fig. S1B-D, Table S4). The genes in PC3 were dominated by epithelial structural and barrier-associated genes, including multiple keratins (*Krt4, Krt5, Krt6a, Krt13, Krt14, Krt15, Krt78*)^46^, cornified envelope components (*Sprr2a2, Sprr2a3, Sprr3, Crct1, Lce3a, Loricrin*)^47^, and desmosomal adhesion proteins (*Dsg1a, Dsg3, Dsc3, Pkp1*)^48^, suggesting that a substantial component of genotype-associated transcriptional variance reflects differences in epithelial barrier integrity between WT and IFNLR1 KO mice. A smaller cluster of skeletal muscle genes was also present among the top PC3 genes.

**Figure 3.**
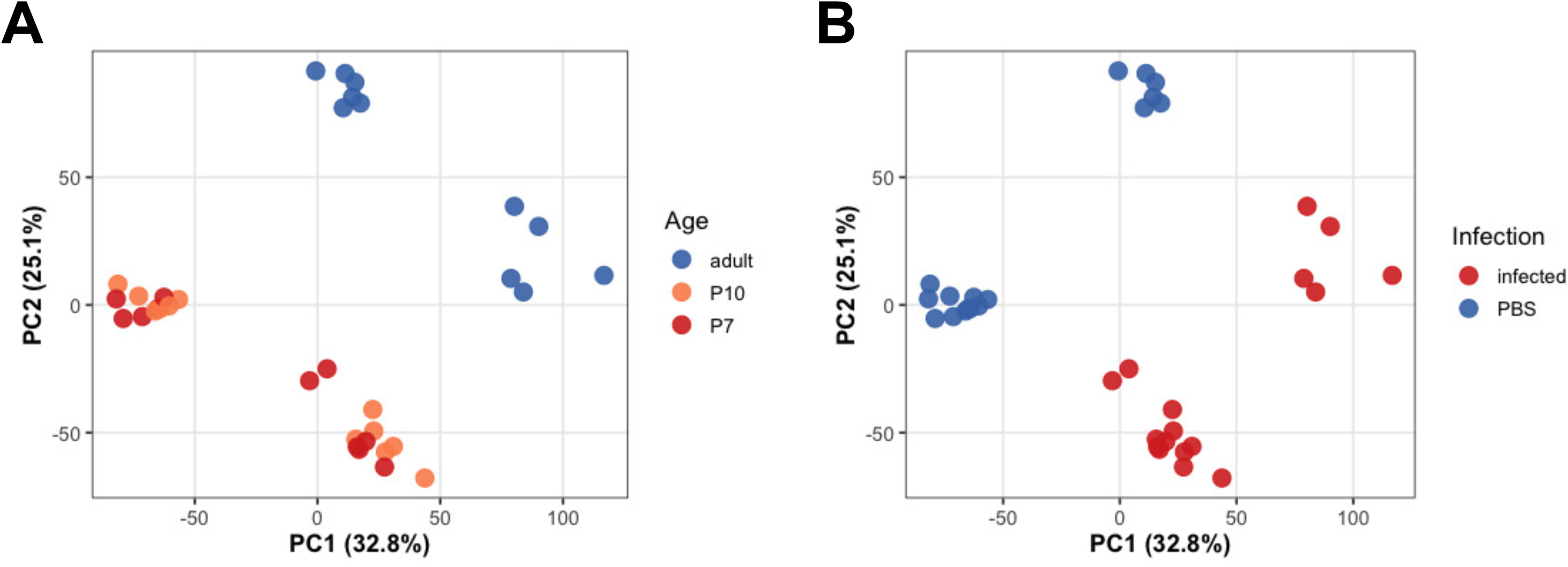
Bulk RNA sequencing reveals age- and infection-dependent transcriptional programs in the lung. Principal component analysis (PCA) of normalized gene expression values from whole lung RNA sequencing of WT and IFNLR1 KO mice inoculated with *B. pertussis* at P7, P10 or adult age and harvested at 7 dpi. Samples are colored by age at inoculation (A) or infection status (B). PC1 accounts for 32.8% of total variance and PC2 accounts for 25.1% of total variance (Fig. S1). n=3 biological replicates per group.

To characterize how *B. pertussis* infection reshapes gene expression across age and genotype, we identified differentially expressed genes (DEGs) in each group by comparing infected mice to their age- and genotype-matched PBS-inoculated controls (Table S1). Based on the survival phenotypes presented in Fig. 1, we strategically selected 4 key groups for transcriptional analysis that best captured the spectrum of infection outcomes: (i) WT P7 and (ii) IFNLR1 KO P10, which both succumb to infection (susceptible groups); (iii) WT P10, which represents the earliest age at which WT animals survive; and (iv) adult WT, which represents fully established resistance. This framework allowed us to directly compare susceptible and resistant phenotypes while bracketing the critical developmental transition from death to survival. Comparing infection-induced DEG sets across these 4 groups revealed both shared and group-specific transcriptional responses (Fig. S2A). A core of 616 upregulated genes (23% of the total upregulated DEG pool across all four groups) was shared across all 4 groups, indicating that a substantial portion of the host transcriptional response to *B. pertussis* is conserved regardless of age or genotype. Pathway enrichment analysis of this core set of genes was dominated by broad, multi-lineage immune terms spanning innate and adaptive immunity (e.g., regulation of immune effector process, leukocyte migration) (data not shown), consistent with general responses to bacterial infection overall. Because IFNLR1 expression in mice is largely restricted to epithelial cells and subsets of immune cells, including neutrophils^19–21,25^, we specifically examined this core set for neutrophil-associated pathway terms. Despite representing a small fraction of the overall enrichment output, four GO Biological Process terms (neutrophil degranulation, regulation of neutrophil degranulation, neutrophil apoptotic process, and neutrophil-mediated killing of symbiont cell) and one KEGG term (neutrophil extracellular trap formation) reached significance, driven by genes including *Cd177*, *Itgam*, *Itgb2*, *Cxcr2*, and *Myd88* (Table S5). This indicates that a conserved neutrophil effector program is embedded within the universal transcriptional response to infection, consistent with the restricted cellular distribution of IFNLR1.

Adult WT mice mounted the most transcriptionally distinct response, with 678 uniquely upregulated genes (25%) (Fig. S2A). In contrast, WT P7 mice induced relatively few unique genes (188, 7%), suggesting that the susceptible infant response is less transcriptionally distinctive and overlaps with the transcriptional response in other groups (Fig. S2A). Pathway enrichment of the 678 genes unique to adult WT infected mice was dominated by cell cycle progression and DNA replication/repair machinery (e.g., chromosome segregation, DNA replication, homologous recombination), co-occurring with genes involved in lymphocyte differentiation and antigen receptor signaling (e.g., *Cd3d*, *Cd28*, *Zap70*, *Foxp3*, *Lag3*) (Table S6). Because *B. pertussis*-specific systemic T cell responses are not typically detected until 2-4 weeks post-infection in naive mice^49^, this proliferative signature at 7 dpi may reflect the larger, more diverse baseline lymphocyte compartment present in mature adult animals. This may include innate-like lymphocyte populations capable of earlier proliferative engagement, rather than *B. pertussis*-antigen-specific clonal expansion, which would not be expected to peak at this early timepoint.

Comparing each P10 genotype to adult mice revealed a notable asymmetry in this shared upregulated signature: WT P10 infected mice shared substantially more genes exclusively with adult WT infected mice (166 genes, 6%) than did IFNLR1 KO P10 mice with adult WT infected mice (54 genes, 2%) (Fig. S2A), raising the possibility that the extent of transcriptional convergence with the mature adult resistance program tracks with protection at P10.

Pathway enrichment analysis of the 166 genes shared exclusively between the two resistant groups (WT P10 and adult WT) was dominated by T cell activation and adaptive immune signaling, including regulation of T cell activation, lymphocyte differentiation, and type II interferon production (Fig 4A and Table S7), driven by genes including *Ifng*, *Cd3e*, *Cd40lg*, *Ctla4*, and *Il2ra* (Table S7). This finding indicates that resistance at P10 is associated not only with IFN-λ signaling but with coordinated engagement of the type II interferon and adaptive T cell compartments (Fig 4A-C and Table S7), consistent with the possibility that resistance depends on interplay between IFN pathways rather than IFN-λ signaling alone.

**Figure 4.**
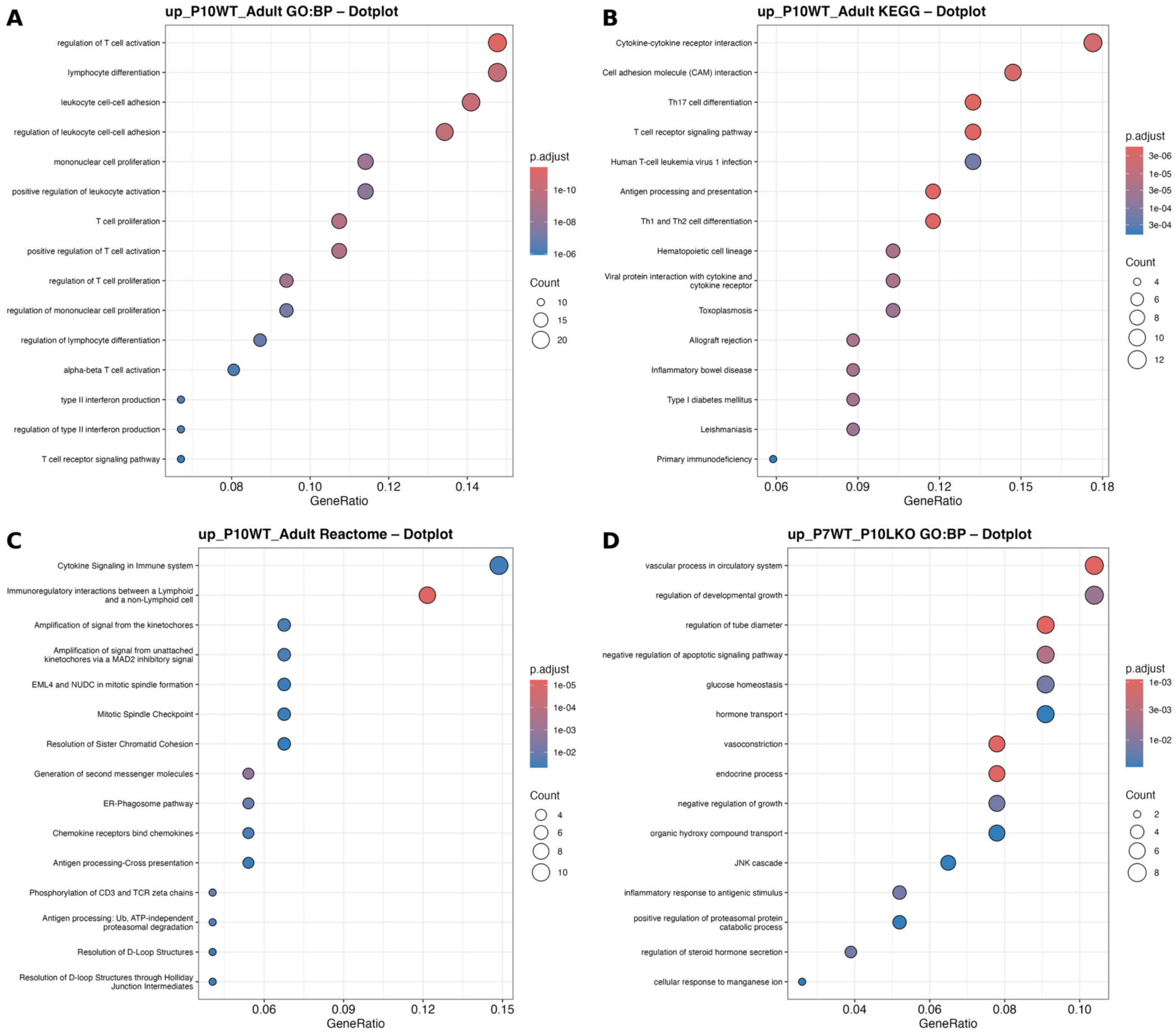
Susceptible and resistant animals engage distinct pathway-level transcriptional programs during *B. pertussis* infection. (A–C) Gene set enrichment dot plots for GO Biological Process (A), KEGG (B), and Reactome (C) databases using the 166 genes shared exclusively between resistant groups (WT P10 and adult WT infected mice) (Table S7). (D) Gene set enrichment dot plot for GO Biological Process using the 83 genes shared exclusively between susceptible groups (WT P7 and IFNLR1 KO P10 infected mice) (Table S8). KEGG and Reactome enrichment of this gene set did not yield significant terms in either database. Dot size represents number of genes contributing to each term. Dot color represents adjusted p-value. All enrichment analyses performed using clusterProfiler with Benjamini-Hochberg correction.

As a complementary analysis, we examined the 83 genes shared exclusively between the two susceptible groups (WT P7 and IFNLR1 KO P10), independent of age or genotype (Fig. S2A). Unexpectedly, the most significantly enriched terms in this comparison were related to vascular tone: vascular process in circulatory system, vasoconstriction, and regulation of tube diameter (Fig 4D and Table S8), driven by genes including *Icam1*, *Alox5*, *Bdkrb2*, and *Hrh1* (Table S8). *Alox5* encodes 5-lipoxygenase, the rate-limiting enzyme in leukotriene biosynthesis, and leukotriene signaling has been directly implicated in pulmonary vasoconstriction and the pathobiology of pulmonary hypertension^50^. This finding parallels a well-described feature of fatal *B. pertussis* infection in human infants: acute pulmonary vasoconstriction and refractory pulmonary hypertension are central, well-documented drivers of mortality in "malignant pertussis" ^51^. This raises the possibility that a lipid-mediator-driven vasoconstrictive program, conserved between infant and IFNLR1-deficient hosts independent of age, contributes to the shared susceptibility phenotype. Lipid transport was also among the enriched terms in this comparison, including *Apol8* (Table S8), consistent with a lipid-handling signature.

A parallel analysis of infection-repressed genes revealed a broadly similar pattern of group-specific and shared responses, though with notable differences in scale (Fig. S2B). A core set of 67 genes (3%) was repressed by infection across all 4 groups. While the immune response to infection appears to follow conserved programs for which genes to induce, repressed genes vary far more dramatically between groups, suggesting that transcriptional suppression is more strongly shaped by the developmental and immunological context of the host than is activation. WT P7 infected mice exhibited by far the largest set of uniquely repressed genes (748, 37%), consistent with a fundamentally distinct baseline transcriptional landscape in the infant lung that renders a large number of genes vulnerable to infection-driven suppression that is not seen in older mice. Adult WT infected mice retained a substantial unique repression signature (347 genes, 17%) (Fig. S2B), though notably smaller in proportion than their unique induced DEG set, suggesting that age-dependent resistance in adults is characterized more strongly by transcriptional activation than suppression. Pathway enrichment analysis was performed on the repressed gene sets shared between susceptible groups (83 genes) and across all four groups (67 genes) (Fig. S2B); neither yielded significant terms in GO Biological Process or Reactome, and only sparse, non-immune-related terms in KEGG (mineral absorption, regulation of lipolysis in adipocytes, cGMP-PKG signaling, cornified envelope formation, protein digestion and absorption) (data not shown). This suggests that, unlike the upregulated response, repressed gene programs do not converge on a shared, biologically coherent pathway across groups. This reinforces the interpretation above that transcriptional suppression is more unique to each developmental and immunological context than is activation.

To directly resolve the transcriptional differences between age-matched animals that differ only in IFN-λ signaling capacity, we performed pairwise differential expression analysis comparing WT P10 infected (resistant) mice and IFNLR1 KO P10 infected (susceptible) mice (Fig. 5A, Table S1). The most significantly upregulated genes in resistant animals included those encoding innate immune effectors, such as IL-22, which promotes epithelial barrier integrity and antimicrobial peptide production at mucosal surfaces^52^. Interestingly, this list also included IFN-γ-inducible genes, such as Ubd, encoding the ubiquitin-like modifier FAT10 that promotes proteasomal degradation of innate immune targets^53^, and *Gbp2b*, a guanylate-binding protein that mediates direct antimicrobial activity against intracellular pathogens^54^. In contrast, susceptible IFNLR1 KO mice showed elevated expression of genes associated with unresolved inflammation and dysregulated lipid handling, *Apol8*, a lipid-interacting apolipoprotein,^55^ and *Selenbp2*, implicated in lipid metabolism and cholesterol efflux^56^. This co-occurrence raises the possibility of interplay between inflammatory and lipid-handling pathways in susceptible animals, though the nature of any such interaction remains to be defined.

**Figure 5.**
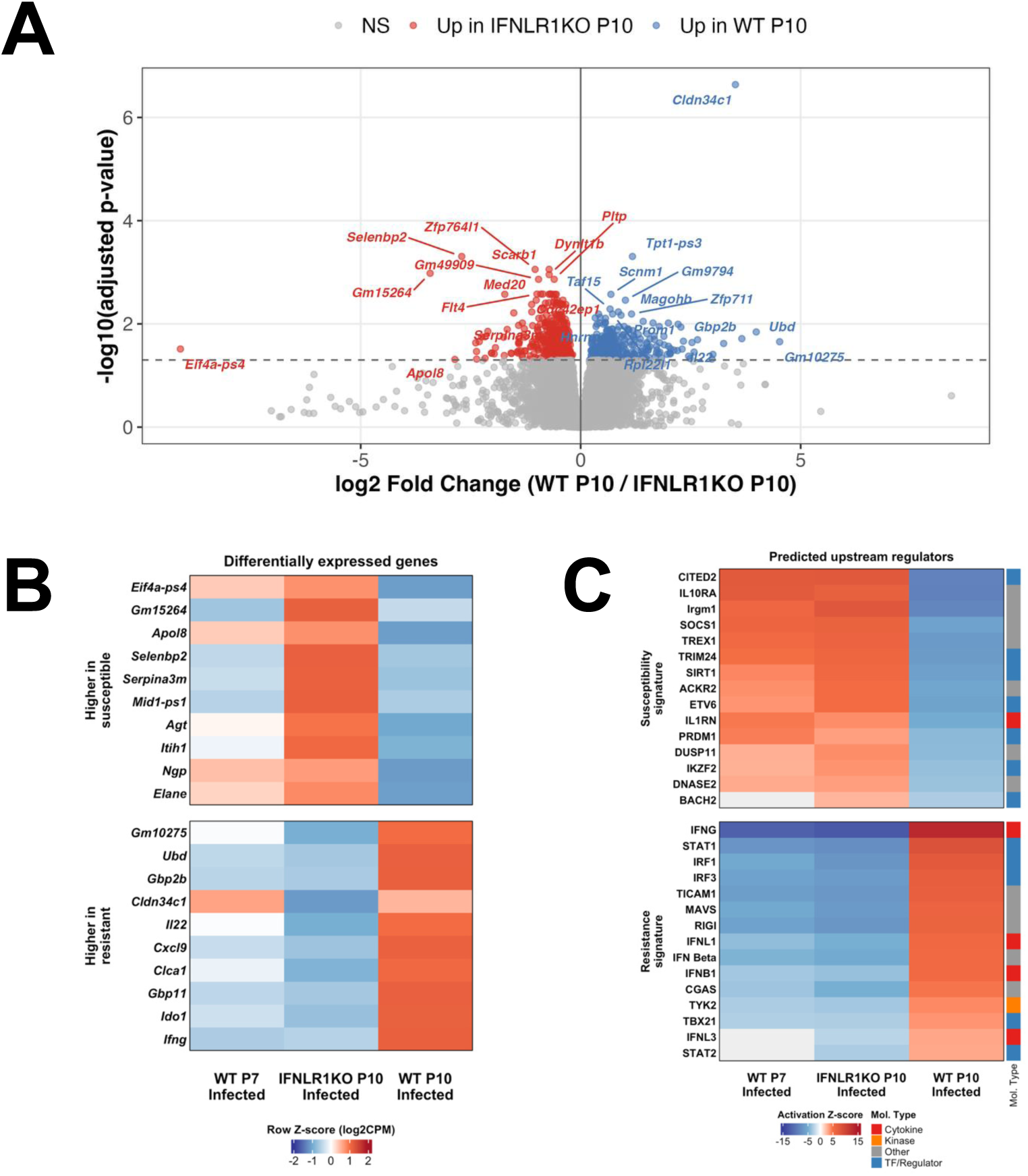
Loss of IFN-λ signaling drives a P7-like transcriptional state in P10 mice during *B. pertussis* infection. (A) Volcano plot of differentially expressed genes (DEGs) at 7 dpi from pairwise comparison of infected WT versus IFNLR1 KO mice inoculated at P10. Genes shown in blue (right) are more highly expressed in WT (resistant) mice; genes shown in red (left) are more highly expressed in IFNLR1 KO (susceptible) mice. The dashed horizontal line represents an adjusted p-value threshold of 0.05 by Benjamini-Hochberg correction. Selected genes of interest are labeled. (B) Heatmap of selected DEGs distinguishing susceptible (WT P7 infected and IFNLR1 KO P10 infected) and resistant (WT P10 infected) groups. Genes displayed were the top 10 genes by log2 fold change among significantly differentially expressed genes (FDR < 0.05, |log2FC| ≥ 1) in infected versus uninfected IFNLR1 KO P10 mice (susceptible signature, top) and the top 10 genes by log2 fold change among significantly differentially expressed genes (FDR < 0.05, |log2FC| ≥ 1) in infected versus uninfected WT P10 mice (resistant signature, bottom). Expression values are displayed as row Z-scores of log2 counts per million (log2CPM), calculated from group mean expression across the three infected groups. Each row is scaled independently so that the Z-score reflects relative expression of that gene across the three groups; values do not represent fold changes relative to uninfected controls. (C) Heatmap of top predicted upstream regulator activation Z-scores from IPA upstream regulator analysis. Regulators are grouped into susceptibility-associated (top) and resistance-associated (bottom) signatures. Molecular type annotations are indicated on the right and below. n=3 biological replicates per group.

The enrichment of both innate immune effector functions and type II IFN-response genes in resistant animals is notable given that only the type III IFN receptor, IFNLR1, was deleted in this model. This also suggests that resistance at P10 may depend on interplay among multiple IFN pathways, rather than IFN-λ signaling acting in isolation.

To visualize expression differences between susceptible (WT P7 and IFNLR1 KO P10) and resistant (WT P10) groups, we constructed a heatmap of the top 10 DEGs by log2 fold change (FDR < 0.05, |log2FC| ≥ 1) in infected versus uninfected IFNLR1 KO P10 mice (susceptible signature) (Fig. 5B upper panel) and the top 10 DEGs by log2 fold change (FDR < 0.05, |log2FC| ≥ 1) in infected versus uninfected WT P10 mice (resistant signature) (Fig. 5B lower panel). IFNLR1 KO P10 infected mice showed an expression pattern across both gene sets that more closely resembled WT P7 infected animals than their age-matched WT P10 counterparts. The susceptible signature included *Apol8* and *Selenbp2*, echoing the lipid-handling genes identified in the shared susceptibility signature described above, and *Agt*, encoding angiotensinogen (Fig. 5B upper panel), the precursor substrate of the renin-angiotensin-aldosterone system^57^. Angiotensin II, the principal proteolytic product of angiotensinogen, is one of the most potent vasoconstrictors known and a central regulator of vascular tone^57^, raising the possibility that this pathway contributes to the same vasoconstrictive phenotype implicated by the *Alox5*-driven leukotriene signature described above (Table S8). The susceptible signature also included *Elane* and *Ngp*, classic azurophilic and specific granule proteins of neutrophils^58^; formal pathway enrichment of the broader shared susceptibility gene set did not support a dominant neutrophil-specific transcriptional program (Table S8), so these two genes are best interpreted as markers of neutrophil granule content rather than evidence of a coordinated neutrophil signature. In contrast, the resistant signature was dominated by *Ifng*, together with *Gbp2b* and *Gbp11*, IFN-γ-inducible antimicrobial GTPases, and *Cxcl9*, an IFN-γ-induced T cell-recruiting chemokine, echoing the type II interferon and T cell activation program identified in the shared resistance signature described above (Table S7). This convergence between WT P7 and IFNLR1 KO P10 expression patterns indicates that loss of IFN-λ signaling does not simply reduce the magnitude of the P10 response but rather shifts IFNLR1 KO P10 mice toward a transcriptional state characteristic of younger, susceptible WT P7 animals.

To identify the upstream regulatory programs driving these differences, we performed Ingenuity Pathway Analysis (IPA)^41^ upstream regulator analysis on DEGs from each of the 3 infected groups (WT P7, IFNLR1 KO P10, and WT P10) by comparing each to their age-matched PBS controls. We then compared the predicted regulator activation scores across the 3 infected groups (Fig. 5C). This comparison revealed two distinct regulatory patterns. WT P7 and IFNLR1 KO P10 infected mice, the 2 groups that succumb to infection, shared predicted activation of regulators associated with immune suppression and attenuation of IFN signaling, including CITED2^59^, SOCS1^60,61^, PRDM1^62^, and BACH2^63^. This signature also included TREX1, a cytosolic DNA exonuclease that limits activation of the cGAS-STING sensing pathway^64^. By contrast, WT P10 infected mice, which survive infection, showed predicted activation of IFN pathway components, including IFNG, STAT1, STAT2, IRF1, IRF3, IFNL1, IFNL3, IFNB1, and TYK2, alongside innate immune sensing regulators TICAM1, RIG-I, MAVS, and cGAS^65^. This resistance signature also included TBX21, encoding the transcription factor T-bet, the master regulator of Th1 differentiation and IFN-γ production^66^. This is consistent with the type II interferon and T cell activation program identified independently in the shared resistance signature described above (Fig 4A-B and Table S7). The fact that WT P7 and IFNLR1 KO P10 animals share a regulatory pattern associated with immune suppression and attenuated DNA-sensing capacity, while WT P10 animals show the opposite pattern of coordinated innate sensing and Th1-driven IFN-γ production, strongly supports the conclusion that IFN-λ signaling is a critical driver of the protective immune response in the lungs that develops between P7 and P10.

### IFNLR1 KO P10 mice exhibit dysregulated pulmonary cellular responses during B. pertussis infection

To test whether the transcriptional patterns described in the previous section were reflected in the underlying cellular composition of the infected lung, we investigated the predicted cellular profile in the lungs across all ages and genotypes represented in the RNA-seq dataset using computational immune deconvolution. We performed deconvolution using 2 methods: (i) mMCP-counter, a marker-based method that estimates the abundance of 16 immune and stromal cell populations from mouse bulk RNA-seq data^42^ (Fig. S3), and (ii) seqImmuCC, an independent algorithm that infers immune cell composition from murine transcriptomic data using a cell-centric gene signature approach^43^ (Fig. S4). Although comparison of infected WT P7, IFNLR1 KO P10 and WT P10 mice showed no differences that reached statistical significance, likely reflecting the limited statistical power of n=3 biological replicates per group, several consistent trends were observed (Fig. 6). mMCP-counter indicated that neutrophil scores were highest in IFNLR1 KO P10 mice (Fig. 6A), although seqImmuCC indicated equivalently high neutrophil scores in WT P7 and IFNLR1 KO P10 mice which were higher than those in WT P10 mice (Fig. 6D), more closely aligning with the survival phenotypes of each group. The elevated neutrophil deconvolution scores in WT P7 and IFNLR1 KO P10 mice indicate that susceptible animals have more neutrophils than resistant animals (Fig. 6A, 6D), and these neutrophils express the same core antibacterial genes identified across all four groups (Table S5). However, this larger number of neutrophils does not translate into better control of infection (Fig 1), suggesting that these neutrophils do not function as effective bacteria-killers in susceptible animals. NK cell deconvolution scores were highest in WT P10 mice, consistent with previous data indicating that higher NK cell abundance associates with improved survival in young mice^4,67–69^ (Fig. 6B, E). T cell scores were also highest in WT P10 mice (Fig. 6C) and seqImmuCC scores indicated that the majority of T cells in young mice are CD8^+^ rather than CD4^+^ T cells (Fig. 6F, G), though this was reversed in adult mice (Fig. S4).

**Figure 6.**
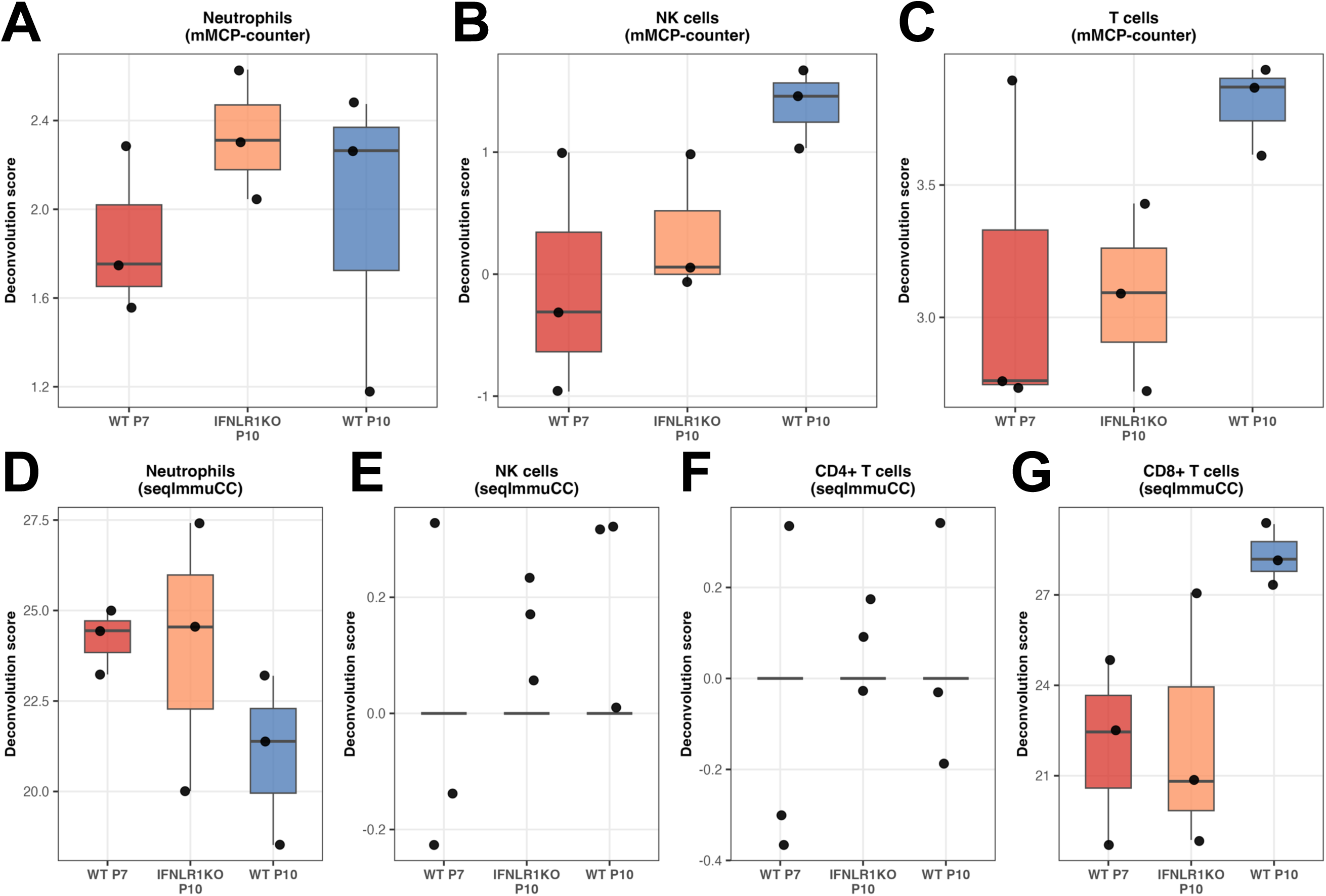
Computational immune deconvolution reveals elevated neutrophil signatures in susceptible mice during *B. pertussis* infection. (A–C) Selected immune cell deconvolution scores estimated by mMCP-counter from bulk RNA-seq data in infected WT P7, IFNLR1 KO P10 and WT P10 mice, showing neutrophil (A), NK cell (B), and T cell (C) populations. (D–G) seqImmuCC was used as an alternative deconvolution method to generate deconvolution scores across the same 3 groups of mice for the following cell types: neutrophils (D), NK cells (E), CD4+ T cells (F) and CD8+ T cells (G). Statistical comparisons performed by unpaired t-test; no individual comparison reached statistical significance. Full deconvolution results across all ages and genotypes are shown in Fig. S3 and Fig S4. n=3 biological replicates per group.

To verify the deconvolution predictions and directly examine pulmonary immune cell populations, we performed flow cytometric analysis of lung immune cells in WT and IFNLR1 KO P10 mice at 7 dpi. IFNLR1 KO P10 mice exhibited significantly elevated numbers of CD45^+^ leukocytes compared to WT P10 mice, after both infection and PBS treatment (Fig. 7A), indicating broadly increased immune cellularity in the lung in the absence of IFN-λ signaling. These differences were not attributable to differences in bacterial burden, as lung bacterial loads were equivalent between genotypes at this timepoint (Fig. 1C). Neutrophil (CD11b^+^Ly6G^+^) counts were more dramatically elevated in IFNLR1 KO P10 infected mice (Fig. 7B), confirming the elevated neutrophil deconvolution scores in susceptible animals (Fig. 6A, D). Because these neutrophils express the same core antibacterial gene program seen in resistant animals (Table S5), yet susceptible animals still fail to control infection, this points to a functional problem in bacterial killing rather than a simple shortage of neutrophils. However, the relative frequency of neutrophils as a proportion of total CD45^+^ cells was comparable between genotypes (data not shown), suggesting that the increase in neutrophil number reflects the overall expansion of pulmonary immune cellularity rather than preferential neutrophil recruitment in the absence of IFN-λ signaling. Resistance of bacteria to neutrophil-mediated killing in susceptible animals may also contribute to the accumulation of these cells in the absence of effective clearance.

**Figure 7.**
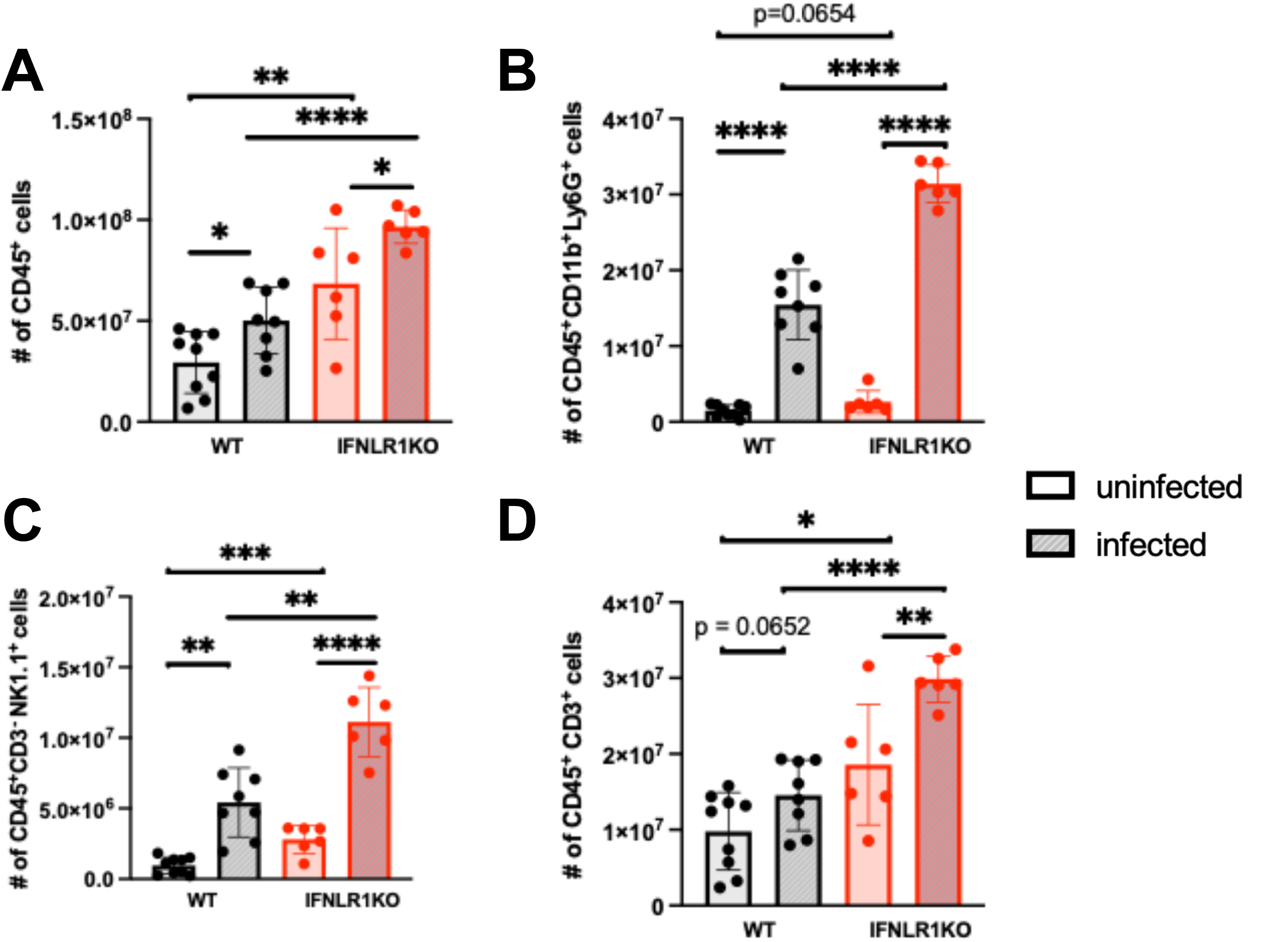
IFNLR1 KO P10 mice exhibit elevated pulmonary immune cellularity during *B. pertussis* infection. Absolute counts of CD45+ leukocytes (A), neutrophils (B), NK cells (C), and CD3+ T cells (D) at 7 dpi in total lung homogenates from WT (black) and IFNLR1 KO (red) mice inoculated at P10 with *B. pertussis* (striped) or PBS (shaded). Each dot represents an individual animal. Statistical comparisons performed by unpaired two-tailed t-test (ns, not significant; *p<0.05; **p<0.01; ***p<0.001; ****p<0.0001; n≥6 per group).

Although not predicted by the deconvolution analysis, NK cell numbers were also higher in IFNLR1 KO mice than in WT P10 mice (Fig. 7C), suggesting that IFN-λ signaling influences NK cell homeostasis. NK cell responses represent a critical determinant of age-dependent protection against *B. pertussis* infection, particularly during the neonatal period when adaptive immunity is immature^4^. The contradictory finding of higher NK cell numbers in susceptible IFNLR1 KO P10 mice than resistant WT P10 mice suggests that IFN-λ signaling may influence NK cell function during infection, although this is likely an indirect effect since NK cells do not express the IFN-λ receptor^70,71^.

Analysis of CD3^+^ T cells also revealed a similar pattern, with higher numbers in IFNLR1 KO mice than in WT P10 mice (Fig. 7D). The accumulation of T cells in IFNLR1 KO P10 infected lungs in the context of increased bacterial dissemination may reflect dysregulated or compensatory recruitment rather than a productive adaptive response, though functional characterization of these cells will be required to fully interpret this finding. Although we did not determine the phenotype of these T cells by flow cytometry, the deconvolution data indicated that the majority of T cells in young mice are CD8^+^ rather than CD4^+^ T cells (Fig. 6F, G). Since CD4^+^ T cells are more protective against *B. pertussis* infection than CD8^+^ cells^49^, these data indicate that lung T cells in these young mice may not play a significant role in protection compared to innate immune cells, although T cell profiles may change later in the infection.

Taken together, the computational deconvolution and flow cytometric data indicate that IFNLR1 KO P10 mice develop broadly elevated pulmonary immune cellularity during infection, with increased numbers of neutrophils, NK cells and T cells. The elevated immune cell signature in susceptible animals, combined with their failure to control bacterial dissemination, suggests that the cellular response in the absence of IFN-λ signaling may be quantitatively expanded but qualitatively impaired, mirroring the broader transcriptional convergence between IFNLR1 KO P10 and WT P7 animals described above (Fig. 5B, C).

## DISCUSSION

In this study, we investigated the role of IFN-λ signaling in shaping age-dependent immune responses to *B. pertussis* infection using a combination of survival analysis, bacterial burden quantification, bulk RNA sequencing, computational immune deconvolution and flow cytometry. Our findings demonstrate that IFN-λ signaling is a critical determinant of the transition from susceptibility to resistance to lethal infection that occurs between P7 and P21 in the infant mouse model of pertussis. Loss of IFNLR1 shifted this protective transition by approximately 10 days, with IFNLR1 KO mice not fully resistant to lethal infection until P21, in contrast to WT mice gaining resistance at P10. IFNLR1 KO P10 mice exhibited enhanced bacterial dissemination and lethality compared to WT age-matched controls. Mechanistically, susceptible P7 WT animals possess a functional IFN-λ receptor in the lungs but fail to produce sufficient IFN-λ during infection for pertussis protection. Consistent with this, supplementation with recombinant IFN-λ prolonged survival in P7 WT animals in an IFNLR1-dependent manner, confirming that restoring IFN-λ signaling can partially overcome infant susceptibility. Transcriptomic analysis revealed that IFNLR1 KO P10 infected mice display a transcriptional state closely resembling WT P7 infected animals rather than their age-matched WT counterparts. This shared susceptibility signature was characterized by suppression of IFN-stimulated genes, innate immune effectors, and adaptive immune priming machinery. Predicted upstream regulators driving this pattern included SOCS1^60,61^, BACH2^63^, CITED2^59^, and PRDM1^62^, all of which are associated with suppression of IFN signaling and immune attenuation. Pathway-level analyses further distinguished susceptible and resistant transcriptional programs, with resistant animals uniquely enriching for antigen processing and presentation, IL-12 family signaling, and IFN-driven effector pathways. Complementary computational deconvolution and flow cytometric analyses revealed dysregulated pulmonary immune cellularity in IFNLR1 KO P10 mice, including elevated neutrophil, NK cell and T cell counts despite the susceptibility of these mice.

The age-dependent insufficiency of IFN-λ production described here is consistent with the broader literature on neonatal IFN deficits^8,72–75^. Neonatal immune cells are well-documented to produce reduced amounts of both type I and type III IFNs in response to innate stimulation, linked to impaired IRF3 and IRF7 activity and epigenetic silencing of IFN loci in neonatal cells^9,10^. Our finding that P7 WT lungs contain functional IFN-λ receptor capable of signaling through STAT1 when stimulated exogenously indicates that this is not the deficient step in infant mice, although we have not yet identified the specific cell type(s) responding to IFN-λ and mediating protection. Instead, the defect is apparently at the level of IFN-λ production in response to infection. This is an important distinction: infant susceptibility in this context is not a fixed developmental limitation of the receptor system, but rather a failure to produce the initiating signal, which in principle could be corrected therapeutically. The ability of exogenous recombinant IFN-λ to prolong survival in P7 WT animals but not in IFNLR1 KO animals confirms that this effect is on-target, consistent with prior reports demonstrating therapeutic potential of IFN-λ supplementation in respiratory mucosal infections^38,39^.

The convergence of IFNLR1 KO P10 and WT P7 infected animals onto a shared susceptibility transcriptional state is among the most striking findings of this study. Rather than simply reducing the magnitude of the P10 immune response, loss of IFN-λ signaling appears to drive a qualitative shift in the character of that response — one that more closely resembles a developmentally earlier state than an attenuated version of the age-appropriate response. Convergent transcriptional susceptibility states between age-disparate groups have been observed in other neonatal bacterial infection models, where impaired IFN responses similarly drive immature-like transcriptional programs regardless of host age^76^. This has implications beyond pertussis: it suggests that IFN-λ signaling at barrier surfaces may be a key driver of immunological maturation of the lung across early postnatal development, and that deficiencies in this pathway could broadly impair the transition from infant to adult-like immune competence. This interpretation is supported by the upstream regulator analysis, which revealed that susceptible animals share predicted activation of SOCS1 and BACH2 — negative regulators of JAK-STAT signaling and IFN responses, respectively. SOCS1 knockout mice develop a fatal neonatal inflammatory disease driven by unrestrained IFN-γ signaling^60,61^, positioning SOCS1 as a critical regulator of the balance between IFN-mediated immunity and immune homeostasis in early life. Its appearance in the shared susceptibility signature of both WT P7 and IFNLR1 KO P10 animals raises the possibility that IFN-λ normally acts to counteract SOCS1-mediated suppression of IFN responses during the critical P10 window.

The parallel analysis of infection-repressed genes revealed that transcriptional suppression during *B. pertussis* infection is considerably more group-specific than transcriptional activation, with only 3% of repressed genes shared across all four groups compared to 11% of induced genes. While the immune response appears to follow broadly conserved programs for which genes to activate during infection, the genes that are repressed vary far more dramatically between groups. One possible explanation is that infection-driven transcriptional suppression is more dependent on chromatin accessibility and the epigenetic landscape of the host cell than activation, both of which differ substantially between neonatal and adult immune cells^9,10^. Neonatal immune cells exhibit broad differences in histone modification patterns, including H3K4me3, and in chromatin accessibility relative to adult cells^9,10^, which may render distinct gene sets susceptible to infection-driven silencing. The disproportionately large unique repression signature in WT P7 animals — 748 genes representing 37% of all repressed DEGs — is consistent with a broadly distinct epigenomic state in neonatal lung tissue that renders a large number of genes susceptible to infection-driven suppression that would not be affected in older animals. Whether this reflects active epigenetic reprogramming by *B. pertussis* virulence factors or a pre-existing feature of the neonatal epigenome remains an important open question.

Among the genes most significantly elevated in resistant WT P10 animals was *Il22*, which encodes a cytokine with well-established roles in epithelial barrier integrity and mucosal antimicrobial defense^28^. IL-22 signals through a receptor expressed predominantly on epithelial cells, promoting production of antimicrobial peptides, mucus, and tight junction proteins that collectively reinforce mucosal barrier function. Notably, both IL-22 and IFN-λ share the IL-10RB receptor subunit and have related roles in respiratory epithelial defense^28^, raising the possibility that IFN-λ signaling promotes IL-22 production or responsiveness as part of a coordinated epithelial defense program. The relationship between IFN-λ and IL-22 at respiratory mucosal surfaces during bacterial infection has not been systematically characterized and represents a compelling direction for future investigation. The role of IFN-λ in respiratory barrier function and biology during *B. pertussis* infection is under active investigation in our lab and will be reported in a separate manuscript.

Pathway analysis of the resistant transcriptional program revealed unique enrichment for IL-12 family signaling^67,68^, a finding that connects directly to our prior work demonstrating a critical role for NK cells and IFN-γ in age-dependent protection against *B. pertussis* infection^4^. IL-12 family cytokines are the primary drivers of IFN-γ production from both NK cells and CD4+ helper T cells and are essential for bridging innate and adaptive immune activation during bacterial infection^67–69^. Their enrichment in resistant but not susceptible animals suggests that effective *B. pertussis* defense requires not only direct IFN signaling through IFNLR1, but also the downstream engagement of IL-12-driven NK cell and Th1 T cell effector programs that collectively amplify local and systemic antimicrobial responses^30,69^. This is further supported by the upstream regulator analysis, which predicted activation of IFNG, TBX21, and STAT1—transcriptional drivers of IFN-γ-producing effector cells—specifically in resistant mice.

The elevated immune cellularity observed in IFNLR1 KO P10 mice, characterized by increased total CD45+ leukocytes and neutrophil, NK cell and T cell subsets, seems counterintuitive since these mice succumb to infection. In the case of neutrophils, this may reflect a broader dysregulation of the pulmonary myeloid compartment rather than selective neutrophil recruitment. One possible explanation is that the cells we identify as neutrophils (CD11b+ Ly6G+) are granulocytic myeloid-derived suppressor cells (PMN-MDSCs). MDSCs are abundantly present in neonates and decline progressively with age^77,78^, a pattern that parallels the age-dependent susceptibility observed in this model. Neonatal PMN-MDSCs share many phenotypic markers and gene expression patterns with neutrophils and suppress both NK cell and T cell responses, including IFN-γ production^78^. During neonatal bacterial infection, MDSCs acquire enhanced immunosuppressive activity, suppressing CD4^+^ T cell proliferation and reducing protective cytokine responses^76^. We have previously shown that *B. pertussis*-infected P7 mice have significantly elevated IL-10 and reduced IFN-γ compared to infected adults^4^, a cytokine profile consistent with MDSC-mediated suppression. Whether IFNLR1 signaling regulates MDSC numbers or function in the neonatal lung during *B. pertussis* infection, and whether the expanded myeloid cellularity in IFNLR1 KO P10 mice reflects MDSC accumulation, represents a compelling direction for future investigation.

The findings of this study have several implications for our understanding of pertussis pathogenesis and for the development of age-targeted therapeutic strategies. The postnatal age window during which murine IFN-λ production becomes established, between P7 and P10, corresponds approximately to the first few weeks of human neonatal life, during which pertussis-related mortality is highest^1,2^. The identification of IFN-λ cytokine insufficiency as a proximal driver of neonatal susceptibility suggests that approaches aimed at augmenting IFN-λ production or signaling in the neonatal lung — whether through adjuvant design, maternal vaccination strategies that prime neonatal mucosal responses, or direct cytokine supplementation — could improve outcomes in the most vulnerable age group. More broadly, the transcriptional framework established here provides a resource for identifying additional molecular targets and biomarkers of severe neonatal pertussis that can be interrogated in future studies. The age-dependent role of IFN-λ described here is likely not unique to *B. pertussis* infection. IFN-λ plays protective roles during respiratory syncytial virus, influenza virus, and SARS-CoV-2 infection at mucosal surfaces^79,80^, and the neonatal deficit in IFN-λ production may contribute to the disproportionate severity of multiple respiratory pathogens in young infants. The systems-level transcriptional framework established here provides a foundation for investigating whether similar IFN-λ–dependent maturation programs operate during other respiratory infections of early life, and whether augmentation of IFN-λ signaling represents a broadly applicable strategy for improving neonatal respiratory immunity.

## ACKNOWLEDGEMENTS

We thank Alicia Bukowski for help with mouse maintenance, Amit Kumar for assistance with some assays and for helpful advice, and Ciaran Skerry, David Rasko, Ashley Mitchell and Emily Flowers for prereviewing the manuscript. Sequencing and some analyses were performed by Maryland Genomics at the Institute for Genome Sciences, University of Maryland School of Medicine. Flow cytometry was performed at the University of Maryland School of Medicine’s & Greenebaum Comprehensive Cancer Center’s Flow Cytometry Core.

Funding for this study was provided by National Institutes of Health grants (to NC) R01 AI141372-S1 and R21 AI168603, a University of Maryland, Baltimore Institute for Clinical and Translational Research grant, and a Howard Hughes Gilliam fellowship award (to DJ). The flow cytometry core was supported by funds through the Maryland Department of Health’s Cigarette Restitution Fund Program and a National Cancer Institute Cancer Center Support Grant (CCSG) P30CA134274.

## Supplemental Figure Legends

**Figure S1.**
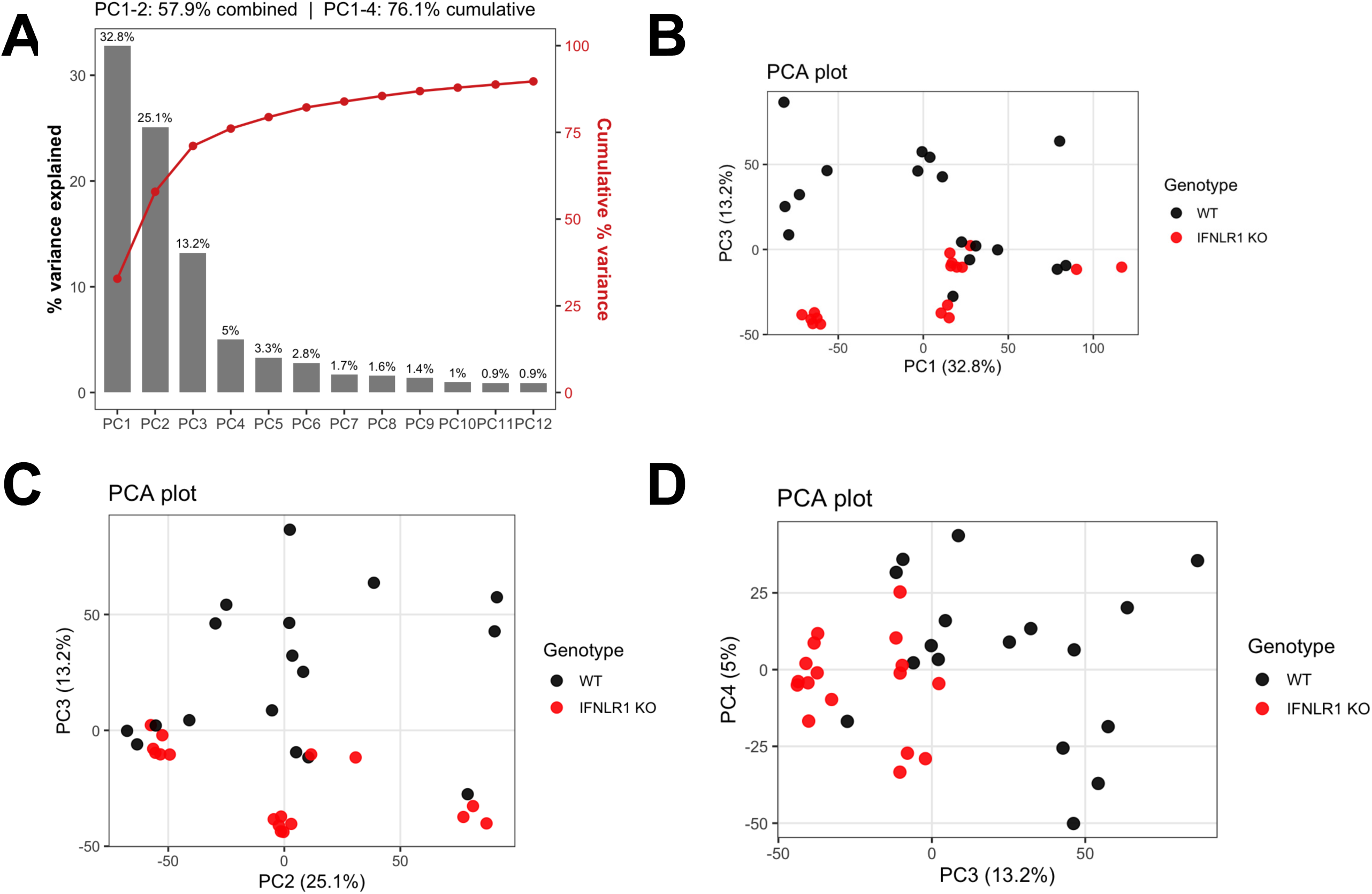
Higher-order principal components reveal a genotype-associated axis of transcriptional variance. (A) Scree plot showing percent variance explained by each of the first 12 principal components from the PCA presented in Fig. 3 (left y-axis, gray bars) and cumulative percent variance indicated (right y-axis, red line and points). PC1 and PC2 together account for 57.9% of total variance; PC1–PC4 account for 76.1% of cumulative variance. (B–D) PCA scatter plots showing PC1 versus PC3 (B), PC2 versus PC3 (C), and PC3 versus PC4 (D), with samples colored by genotype (WT, black; IFNLR1 KO, red). Although genotype is not well resolved along PC1 or PC2 (Fig. 3), PC3 partially separates samples by genotype, suggesting that a genotype-associated axis of transcriptional variance exists within the dataset beyond the age- and infection-associated variance captured by the first two components. n=3 biological replicates per group.

**Figure S2.**
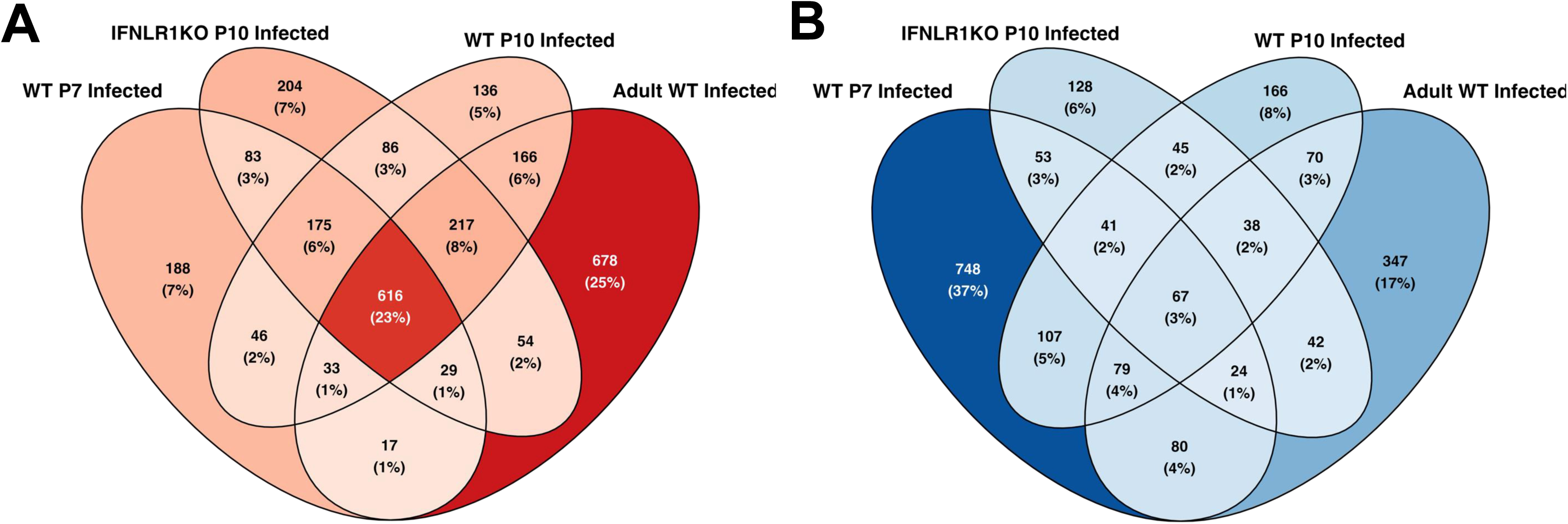
DEG overlap across susceptible and resistant groups. (A) Four-way Venn diagram showing overlap of infection-induced DEGs (FDR ≤ 0.05, |log2FC| ≥ 1) identified by comparing *B. pertussis*-infected mice to age- and genotype-matched PBS controls in WT P7, WT P10, IFNLR1KO P10 and WT adult groups. Each number indicates the count of genes unique to or shared between the indicated groups. Percentages reflect the proportion of that gene count out of the total pool of unique DEGs identified across all four groups combined. (B) Four-way Venn diagram showing overlap of infection-repressed DEGs (FDR ≤ 0.05, |log2FC| ≥ 1) across the same four groups and comparisons as in (A). Numbers and percentages are calculated as described for panel (A). n=3 biological replicates per group.

**Figure S3.**
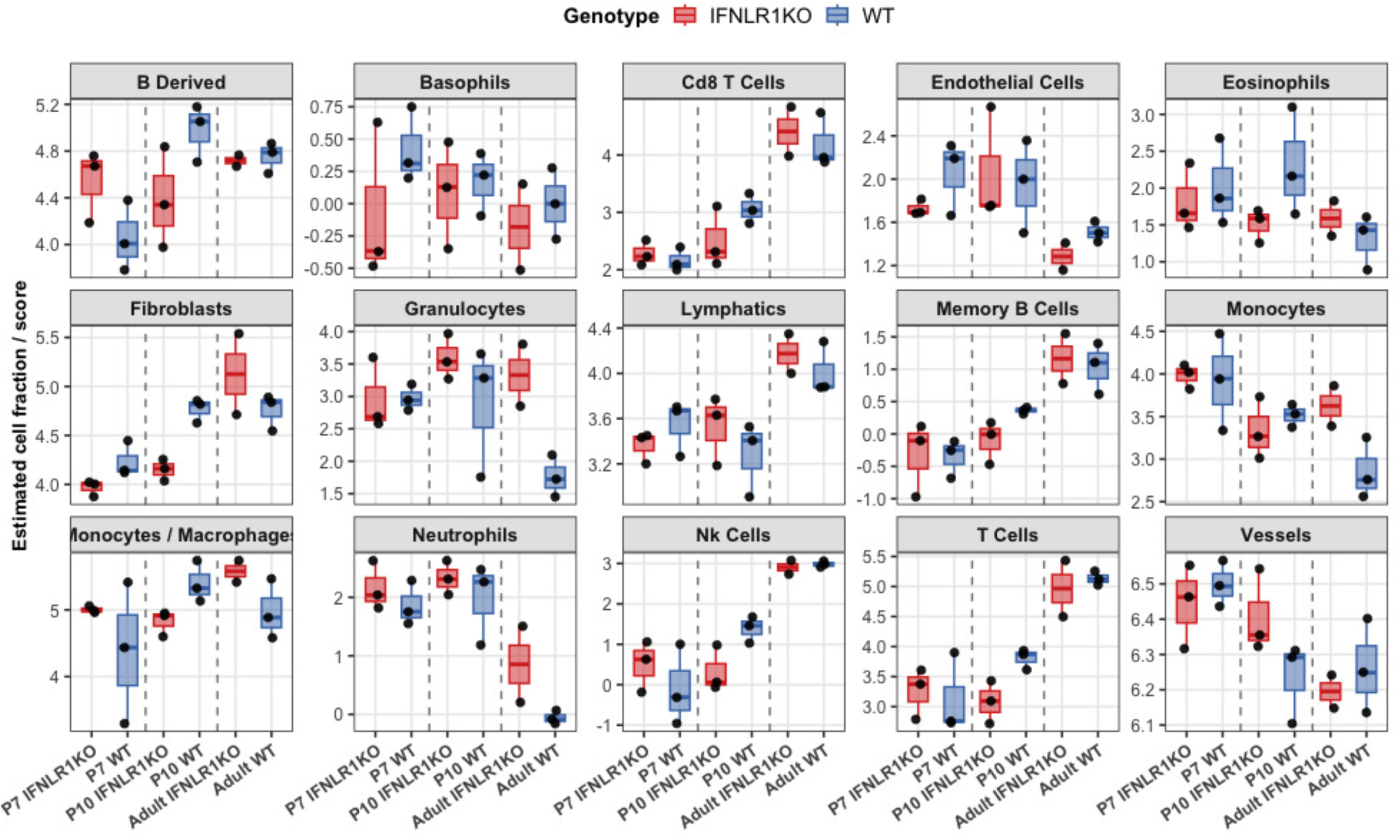
Full mMCP-counter immune cell deconvolution across all ages and genotypes during *B. pertussis* infection. Immune cell deconvolution scores estimated by mMCP-counter from bulk RNA-seq data across all infected age and genotype groups, showing all 15 cell population scores. Data are presented as exploratory trends supporting the focused comparisons shown in Fig. 6. n=3 biological replicates per group.

**Figure S4.**
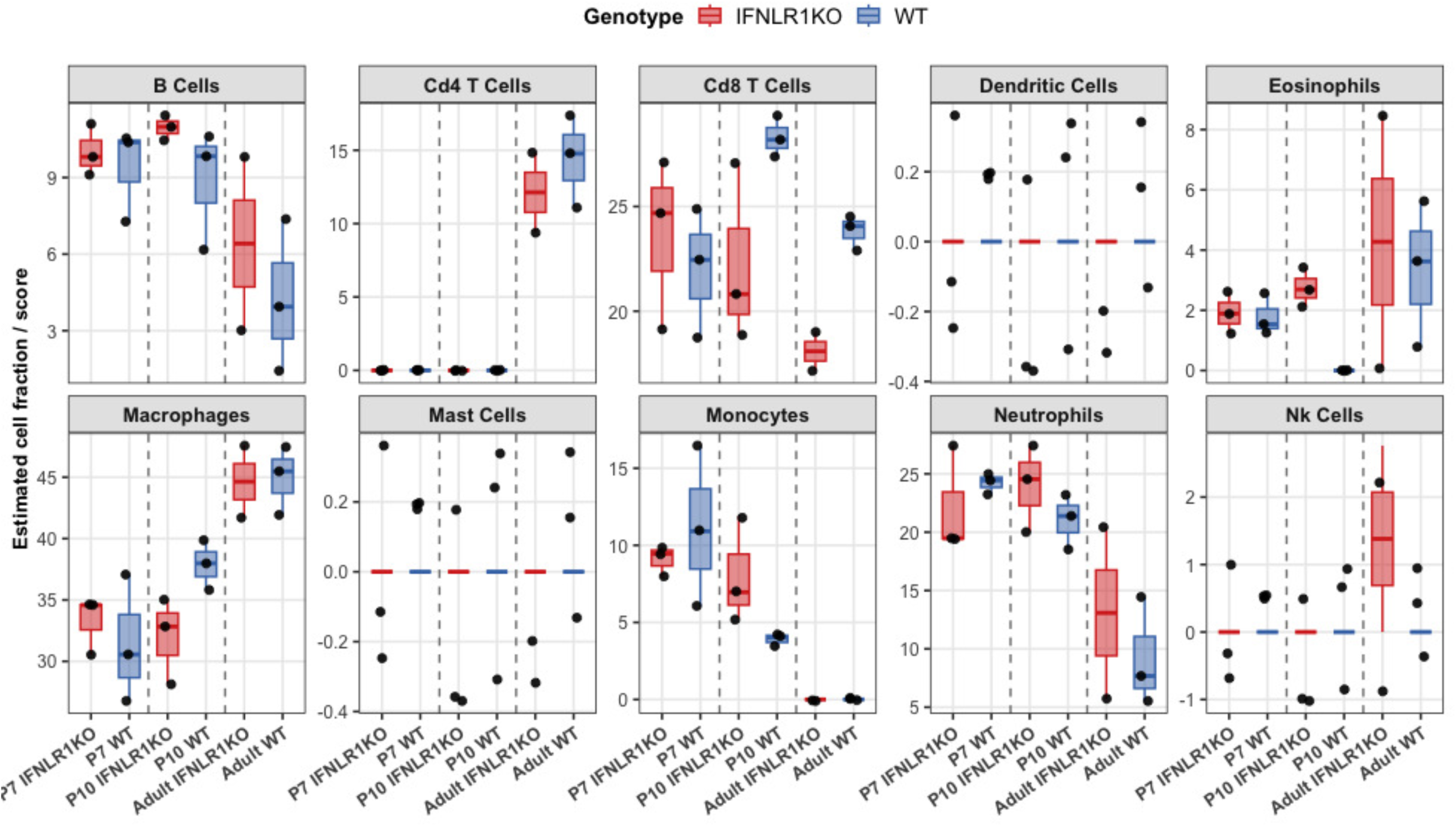
Full seqImmuCC immune cell deconvolution across all ages and genotypes during *B. pertussis* infection. Immune cell deconvolution estimates from seqImmuCC across all infected age and genotype groups, showing all 10 immune cell population scores. Data are presented as exploratory trends supporting the focused comparisons shown in Fig. 6. n=3 biological replicates per group.

